# Renegade Bacterial Genetic Sequences in a Stealth Adapted Virus: Biological and Diagnostic Implications

**DOI:** 10.1101/2022.10.11.511846

**Authors:** W. John Martin

## Abstract

There are major differences between viruses, bacteria, and eukaryotic cells in the structuring of their genomes, modes of replication, and capacity to horizontally transfer genetic sequences. DNA sequencing studies on a virus cultured from a patient with chronic fatigue syndrome (CFS) have confirmed a previously underappreciated capacity of certain viruses to capture and transfer bacterial and cellular genetic sequences between eukaryotic cells as part of the infectious process. The virus originated from an African green monkey simian cytomegalovirus (SCMV). It is termed a stealth adapted virus since infection is not accompanied by inflammation. The immune evasion is attributed to the loss and mutation of the genes coding for the relatively few components that are normally targeted by the cellular immune system. This article provides further elucidation of the origins of many of the bacterial-derived genetic sequences present in the virus. There are multiple clones with close but non-identical sequence alignments with different genomic regions of the *Ochrobactrum quorumnocens* A44 species of bacteria. Another set of clones matched most closely to diverse genomic regions of *Mycoplasma fermentans* bacteria. The sequences of several other clones could only be approximately aligned to those of different types of bacteria. The sequence of clone 3B513 is consistent with genetic contributions from the genomes of several types of bacteria. The term viteria refers to viruses with bacteria-derived genetic sequences. They are the likely primary cause of CFS and autism, and to act as major cofactors in many illnesses, including AIDS. As a more general phenomenon viteria with different types of renegade bacterial sequences can lead to the mistaken diagnoses of bacterial rather than viral diseases. It is important to genetically sequence additional stealth adapted viruses from patients with a wide range of illnesses, including those currently being attributed to Mycoplasma, Borrelia, or Streptococcal infections.

## Introduction

Molecular analysis of cloned DNA derived from viral cultures from a patient with chronic fatigue syndrome (CFS) showed that the cultured virus had originated from an African green monkey simian cytomegalovirus (SCMV) [1–4]. Yet, genetic sequences corresponding to major regions of the SCMV genome were not detected within any of the sequenced DNA clones [4–5]. Moreover, there was an uneven distribution of the clones with regards to the remaining identified regions of SCMV, with genetic variability within clones that match to the same region of the SCMV genome. These findings are consistent with an immune escape mechanism, referred to as stealth adaptation, occurring from the deletion or mutation of the genes coding the relatively few virus components that are normally targeted by the cellular immune system [1, 6]. In addition to the SCMV-derived genetic sequences, there are certain clones with genetic sequences that have come from portions of cellular and bacterial genomes [7–13]. These incorporated cellular and bacterial renegade sequences may be required for the virus to regain its infectivity and may also contribute to the virus-mediated cytopathic effect (CPE).

The increasing availability of DNA sequence data in GenBank has allowed further elucidation of the origins of bacterial-derived genetic sequences present in this virus. The findings are consistent with genetic contributions to the virus genome not only from intracellular growing bacteria, such as *Mycoplasma fermentans* [14], but also from soil-based bacteria including members of the *Brucella – Ochrobactrum* Family within the *Hyphomicrobiales* Order [15]. The findings have major biological implications and are relevant to the potential of misdiagnosing stealth adapted virus infections as being caused by bacteria with patients being inappropriately treated with antibiotics.

## Materials and Methods

Patient and Virus Culture: The virus was repeatedly cultured from a woman who was hospitalized in 1990 with a provisional diagnosis of either encephalitis or meningitis developing a week after she had a sore throat [1]. A cerebrospinal fluid (CSF) sample obtained during her hospital admission showed no cells and a normal protein level. She failed to regain her prior level of health and vigor. She felt cognitively impaired, constantly fatigued, and lost the capacity for restorative sleep. She had earlier rented an apartment to a HIV positive individual with similar symptoms and considered the possibility of a non-sexually transmitted illness. The woman provided a blood sample as part of a study using the polymerase chain reaction (PCR) to look for viruses in patients with CFS. Based on a clearly positive PCR result, an additional blood sample was obtained for virus culturing. A cytopathic effect (CPE) was observed in primary human foreskin fibroblasts (MRHF). It comprised the formation of foamy vacuolated cells with marked syncytia [1]. Subsequently obtained blood samples confirmed the presence of an infectious cytopathic agent that could be passaged in cultured cells. Abundant viral particles were seen by electron microscopy. The cultured cells and cell free supernatants yielded strongly positive PCR assays with well-defined PCR products shown in agarose gel electrophoresis. Radiolabeled PCR products hybridized with material that was pelleted by ultracentifugation of 0.45 μ filtered culture supernatant. As was shown in the earlier publication and included in the present article, DNA extracted from this material migrated in agarose gel electrophoresis as a well-defined band with an estimated size of approximately 20 kilobase (kb).

Cloning of Virus DNA: In the first set of cloning experiments, the DNA obtained from the filtered and ultracentrifuged culture supernatant was digested overnight with 10U/μl of EcoRI enzyme and cloned into pBluescript plasmids [1]. The plasmids were propagated in XL-1 bacteria and those with inserts were either partially or fully sequenced. This series of over 180 clones was labeled as the 3B series. In a later cloning experiment, the nucleic acids extracted from the filtered and ultracentrifuged culture supernatant were further purified by agarose gel electrophoresis. The approximately 20 kb band was excised, and the DNA was reextracted, and then digested overnight with 2U/μl SacI restriction enzyme [5]. The digested DNA was cloned into pBluescript plasmids and propagated in XL-1 bacteria. The resulting series of over 120 clones was labeled as the C16 series. The individual clones of stealth virus-1 that are referred to in this article, together with their NCBI Accession number and nucleotide length are listed in Table 1. Separate Accession numbers corresponding to the separate sequences derived from T3 and T7 polymerase primer sites of the pBluescript plasmids are provided for those clones that were not completely sequenced.

**Table 1.**
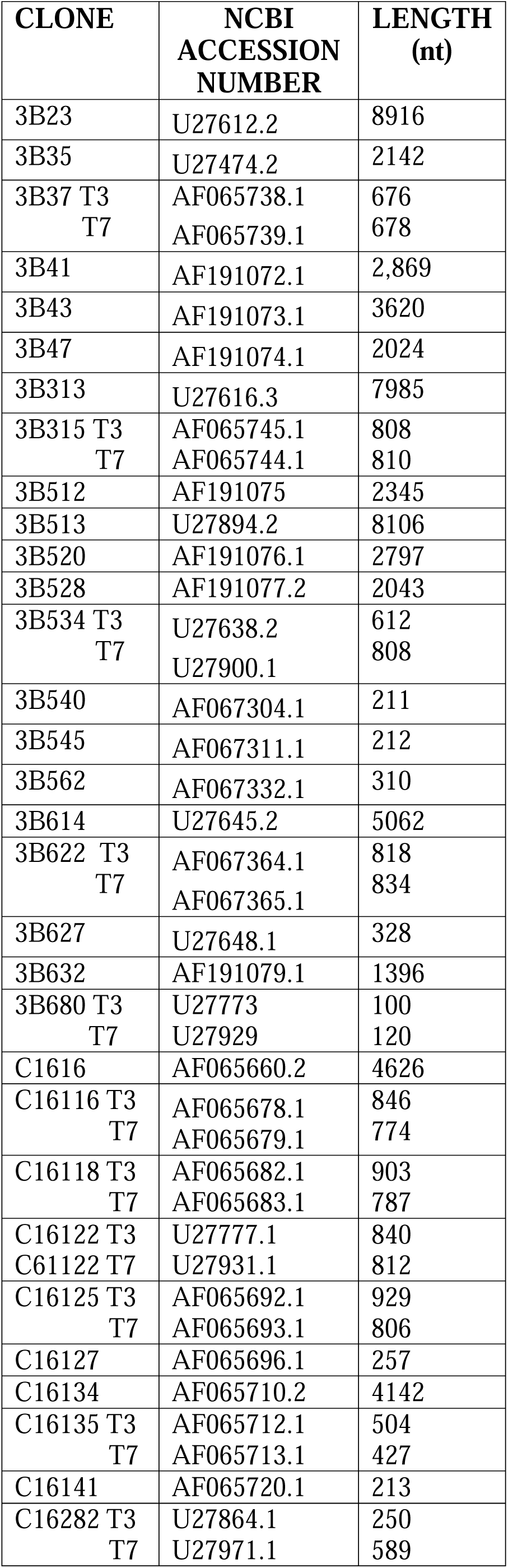
Designations and NCBI Accession Numbers of Clones of Stealth Virus-1 with Matching Bacterial Sequences.

| **CLONE** | **NCBI**<br>**ACCESSION**<br>**NUMBER** | **LENGTH**<br>**(nt)** |
| --- | --- | --- |
| 3B23 | U27612.2 | 8916 |
| 3B35 | U27474.2 | 2142 |
| 3B37 T3<br>T7 | AF065738.1<br>AF065739.1 | 676<br>678 |
| 3B41 | AF191072.1 | 2,869 |
| 3B43 | AF191073.1 | 3620 |
| 3B47 | AF191074.1 | 2024 |
| 3B313 | U27616.3 | 7985 |
| 3B315 T3<br>T7 | AF065745.1<br>AF065744.1 | 808<br>810 |
| 3B512 | AF191075 | 2345 |
| 3B513 | U27894.2 | 8106 |
| 3B520 | AF191076.1 | 2797 |
| 3B528 | AF191077.2 | 2043 |
| 3B534 T3<br>T7 | U27638.2<br>U27900.1 | 612<br>808 |
| 3B540 | AF067304.1 | 211 |
| 3B545 | AF067311.1 | 212 |
| 3B562 | AF067332.1 | 310 |
| 3B614 | U27645.2 | 5062 |
| 3B622 T3<br>T7 | AF067364.1<br>AF067365.1 | 818<br>834 |
| 3B627 | U27648.1 | 328 |
| 3B632 | AF191079.1 | 1396 |
| 3B680 T3<br>T7 | U27773<br>U27929 | 100<br>120 |
| C1616 | AF065660.2 | 4626 |
| C16116 T3<br>T7 | AF065678.1<br>AF065679.1 | 846<br>774 |
| C16118 T3<br>T7 | AF065682.1<br>AF065683.1 | 903<br>787 |
| C16122 T3 | U27777.1 | 840 |
| C61122 T7 | U27931.1 | 812 |
| C16125 T3<br>T7 | AF065692.1<br>AF065693.1 | 929<br>806 |
| C16127 | AF065696.1 | 257 |
| C16134 | AF065710.2 | 4142 |
| C16135 T3<br>T7 | AF065712.1<br>AF065713.1 | 504<br>427 |
| C16141 | AF065720.1 | 213 |
| C16282 T3<br>T7 | U27864.1<br>U27971.1 | 250<br>589 |

Sequencing of Clones: Preliminary sequencing of the clones was performed from both the T3 and the T7 primer sites. The initial sequencing was performed by either Bio Serve Biotechnologies, Laural MD or by the City of Hope Cancer Center Molecular Core Facility, Duarte CA. Sequenase was used at the first location and generally yielded unambiguous 100-200 nucleotide sequences. Thermal cycling was used at the second facility and would commonly yield over 500 nucleotides, but with increasing numbers of undefined nucleotides indicated as “N’; in the longer stretches. Only the T3 and T7 sequences are available for many of the entire 3B and C16 series of clones. Additional double stranded complete sequencing was, however, obtained for most of the clones discussed in this article. The extended sequencing was provided by Lark Technology, Houston, Texas and by U.S. Biochemical Corp. Cleveland, Ohio. All DNA sequence data on the clones have been submitted to GenBank under the heading of stealth virus-1.

DNA Sequence Analysis: The sequences of the clones were analyzed using the online BLASTN and BLASTX programs provided by the National Center for Biotechnology Information (NCBI) [16]. The BLASTN sequence matching and alignment program compares the input (“query”) DNA sequence with the nt/nr non-redundant DNA “subject” sequences currently available in the GenBank repository. The BLASTX program compares the sequences of each of the six potentially derived amino acid sequences coded by the query input sequence with the known protein subject sequences on GenBank. When an amino acid match is identified, the coding nucleotide sequence of the known protein can be used in a pairwise BLASTN comparison with the query sequence. The designated name, NCBI accession number, and the range of nucleotide numbers in the identified GenBank subject sequences are displayed in the BLASTN results. If the input sequence is already in GenBank, this sequence will be the first selected sequence, followed by the sequences with increasing nucleotide disparities till the set limit of identified matching sequences is met. The BLASTN program further tests if the placing of one or more gaps in the input and selected sequences, or if separating the sequences into two or more segments, will improve the overall alignment. The “bit score” is a measure of the non-random chance alignment of two sequences. The number of inserted gaps to obtain optimal alignment is also shown in the results, along with the numbers and ratio of identical nucleotides. The statistical probability against a random alignment is more clearly reflected in the Expect Value. This is expressed as the negative exponential value to the log base “e.” The smaller the “e” value, i.e., the higher the negative log value, then the less likelihood of purely random matching. When the e value is less than e-180, it is recorded as 0.0. This reflects that the sequences are very similar over sufficiently long regions. There can be multiple GenBank sequences that show close matching to only a portion of the query sequence. This can potentially reduce the sensitivity of showing less well aligned sequences. To avoid this, the BLASTN analysis was typically repeated after excluding the more highly matched regions.

## Results

A previously published Figure is included in this article as Figure 1. It shows an ethidium bromide-stained agarose gel with HindIII and BetE-II enzyme digested Lambda DNA phage size markers in the left and right outside lanes, respectively. The added arrow indicates the clearly seen band of a portion of the DNA extracted from the material that was pelleted by ultracentrifugation of filtered cultured supernatant. In comparison with the DNA size markers, the band has DNA of approximately 20 kb. The lane directly beneath the arrow shows several smaller DNA bands, which resulted from EcoRI digestion of another portion of the pelleted and extracted DNA. The lower stained material in the lanes of the supernatant extracted material is RNA. The remaining lane is an EcoR1 digest of nucleic acids extracted from lysed infected cells of the culture from which the supernatant was obtained.

**Figure 1.**
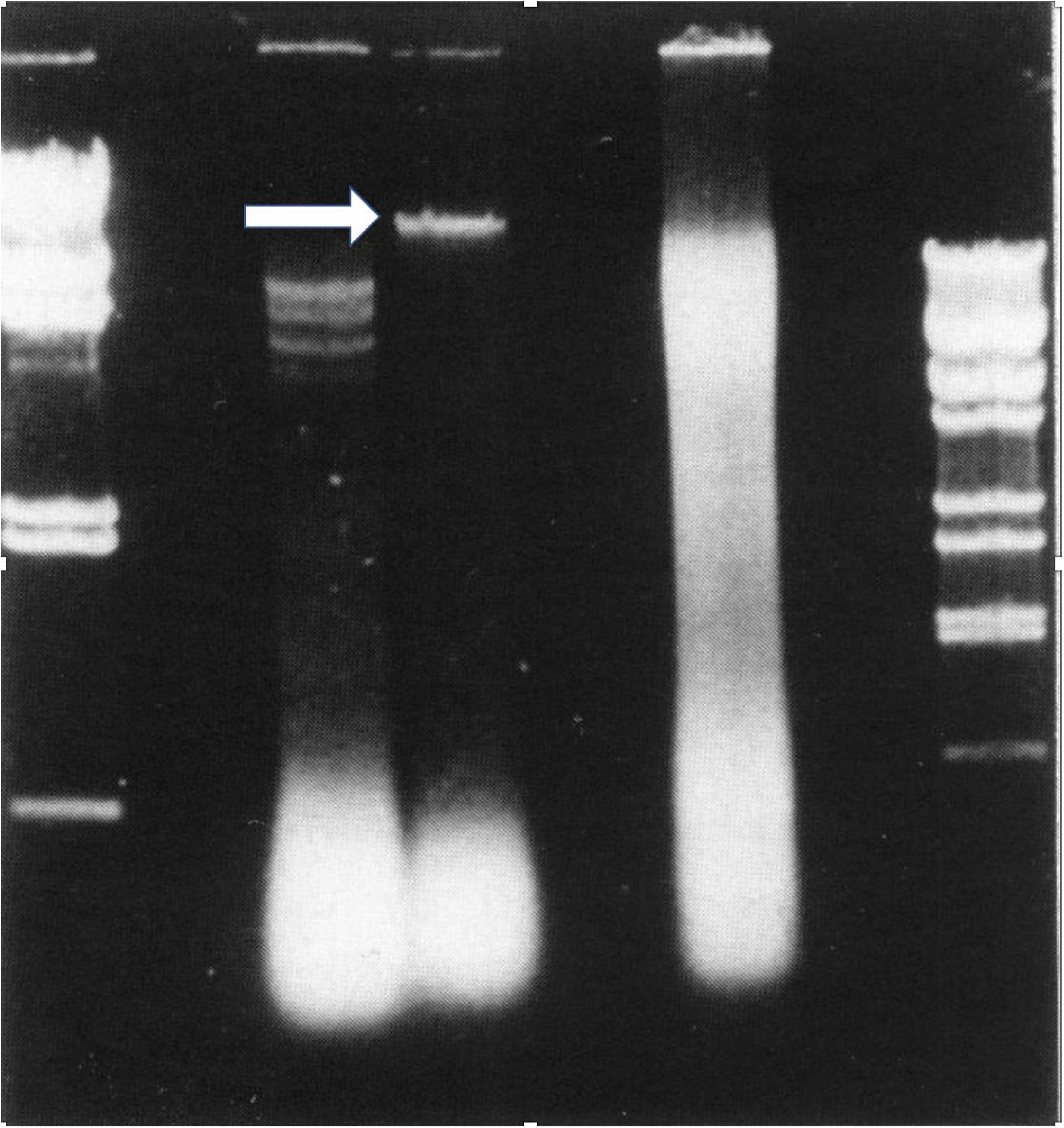
Ethidium-stained agarose gel of electrophorized nucleic acids extracted from the pelleted materials filtered from ultracentrifuged culture supernatant of virus infected MRHF cells (indicated by the arrow) and from the lysate of frozen-thawed infected MRHF cells. The whole cell lysate and a portion of the pelleted material obtained from the culture supernatant were both digested with EcoRI restriction enzyme. The lane to the right of the arrow is the EcoRI digest of the whole cell lysate. The lane beneath the arrow is the EcoRI digest of the pelleted material. The material staining in the lower regions of the internal lanes is RNA. The left and right-side outer lanes are HindIII and BstE-II cut lambda phage. The largest DNA markers for these enzymes are 23,130 bp and 8,454 bp, respectively. The size of the band of uncut DNA is approximately 20 kb. Note also that except at the point of application, there is no discernable DNA staining above the 20 kb band. The Figure is from reference 1.

### Clones With Sequences Matching to *Brucella* - *Ochrobactrum* Bacteria

BLASTN analyses of the DNA sequences in eight of the clones show that the highest levels of non-self-matching to DNA sequences are to distinct regions of Chromosome 1 of *Ochrobactrum quorumnocens* strain A44 [17; NCBI accession number CP022604.1]. The clones are listed in Table 2 in the order of the regions of the matching nucleotides along chromosome 1 of the bacterium. These regions code for various functional proteins, which will be detailed in a future publication. Each of the eight clones had been fully sequenced. The overall ratio of identical nucleotides (nt) in the eight clones to the matched regions in chromosome 1 of *O. quorumnocens* strain A44 is 99.7% (31,977/32,064 nt). Except for the 11 gaps for clone C16116, there is on average only 1 gap per clone for the other 7 clones.

**Table 2.**
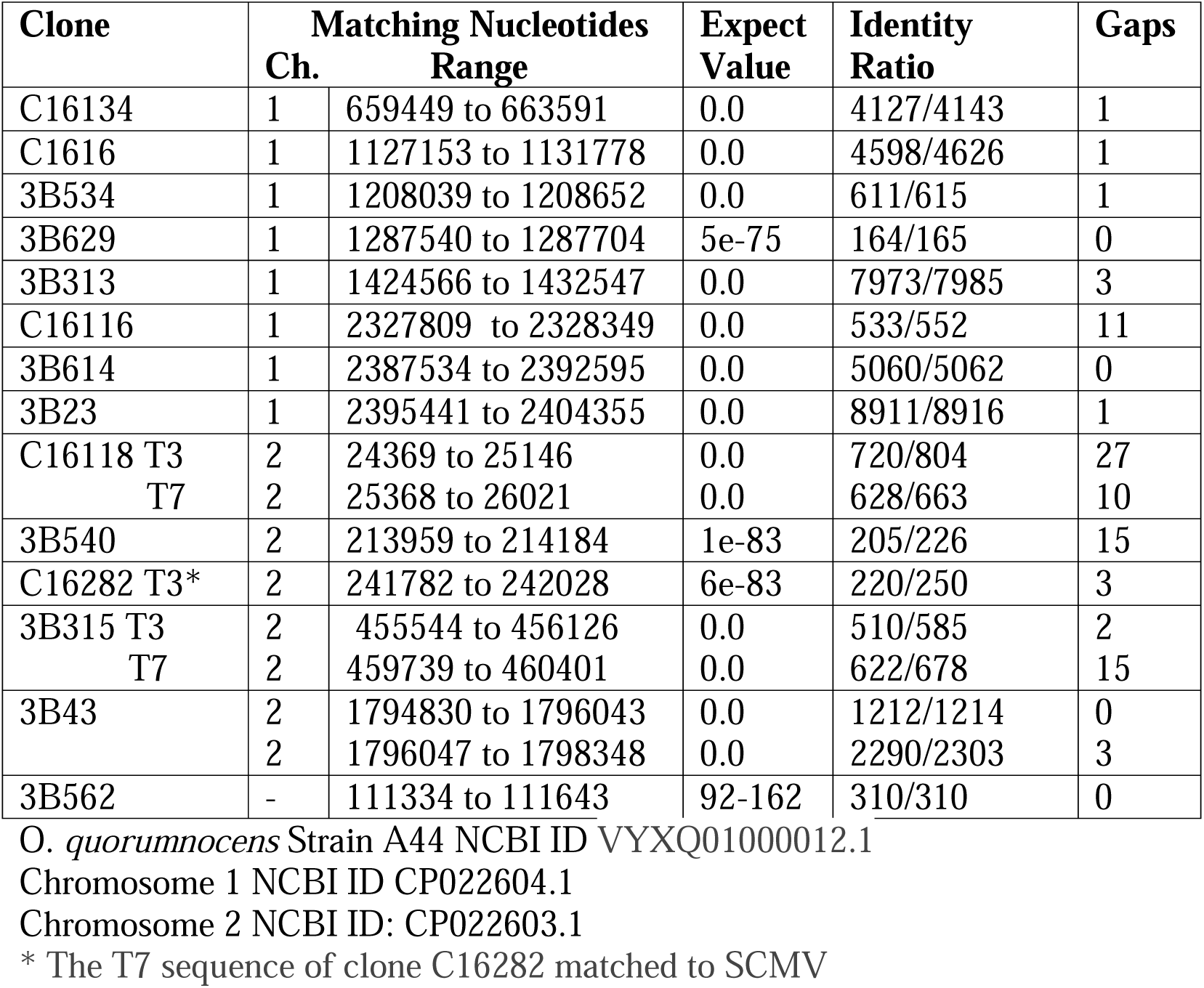
Clones With Sequences That Best Match to *Ochrobactrum quorumnocens* Strain A44.

There are three other clones that were only partially sequenced from the T3 and T7 promoter sites on the pBluescript plasmid. For two of these clones (C16118 and 3B315) both ends of the clones matched to regions within Chromosome 2 of *O. quorumnocens* strain A44 (NCBI accession number <u>CP022603.1)</u>. The T3-derived sequence for plasmid C16282 also matched to a region of chromosome 2 of *O. quorumnocens* strain A44. The T7-derived sequence, however, matched to a genomic region of SCMV. The matching to SCMV strain 2715 (NCBI accession number FJ483968.2) extended from nucleotide 84420 and 84692 with an Expect Value of 1e-63, and 192/248 identical nucleotides after excluding unmatchable “N” nucleotides in the clone. A small fully sequenced clone (3B540) also matched preferable to a region in chromosome 2 of *O. quorumnocens strain A44*. There were 205/226 identical nucleotides, although the optimal matching required 15 gaps.

There is an additional fully sequenced clone (3B43) with 3620 nucleotides. BLASTN identified the statistically higher overall match to the ribosomal genetic sequences in *Brucella BTU1*, *Brucella pituitosa* strain AA2, and *Brucella pseudogrignonensis* strains . Note that the term *Brucella* in referring to these later two strains is a synomon for *Ochrobactrum*. There was less matching to *O. quorumnocens* strain A44 genome. The lower matching occurred because the BLASTN program matched the overall sequence of clone 3B43 to the *Brucella* species, but to two separated ribosomal sequences in the *O. quorumnocens* strain A44 genome. A 103 nucleotide long stretch of the 3B43 clone from nucleotide 2301 to nucleotide 2404 was present at a 99-100% identity in the earlier mentioned *Brucella* strains, but not in the O. quorumnocens strain A44. The missing sequence corresponds to part of the ribosomal gene complex that intervenes between the 16S and the 28S coding regions. Intervening nucleotides are present in some but not of the bacteria within the different bacterial families [18]. Sequences in the two regions of clone 3B43 that are apart from the intervening region match most closely to those in Chromosome 2 of *O. quorumnocens* strain A44. These sequences were, therefore, chosen to be included in of Table 2.

The initial BLASTN results with clone 3B562 did not include any *Onchrobactrum* species. Rather, it identified a sequence in *Serratia marcescens* bacteria with 88% nucleotide identity. Yet, the highest matching to its BLASTX translated amino acid sequence was to an *O. quorumnocens* coded protein. The 310-nucleotide sequence coding the matching region of the *O. quorumnocens* protein is identical to the sequence of clone 3B562. For convenience, this entity has been added to Table 2, even though it could not be specifically identified in the published complete sequences of either chromosome 1 or 2 of *O. quorumnocens*.

There are sequences in two clones, 3B41 and 3B47, that do not closely align with sequences in either chromosome 1 or 2 of *O. quorumnocens strain A44* genome but do align better to sequences in chromosome 2 of *Brucella pseudogrignonensis* [19]. Even though the Expect Value was still 0.0, the total identical matching of the two clones was only 77.1% (3779/4901) with a total of 82 gaps (Table 3).

**Table 3.**
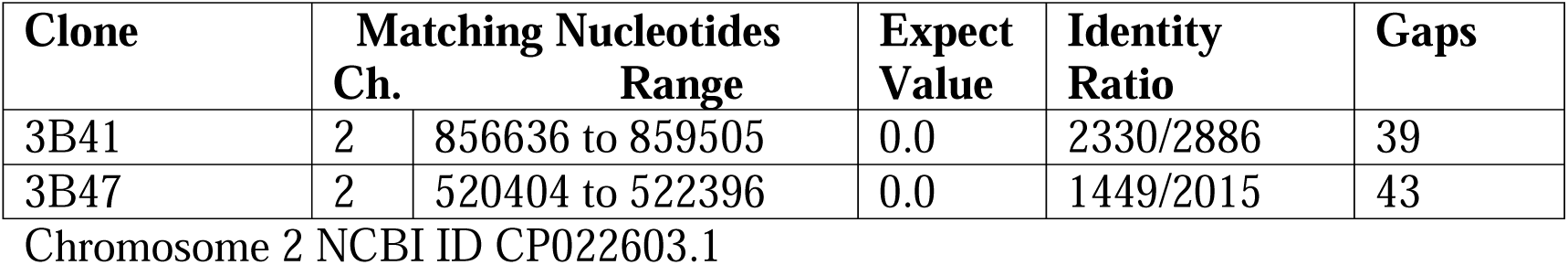
Clones With Sequences That Best Match to *Brucella Pseudogrignonensis* Strain K8.

| Clone | Matching Nucleotides |  | Expect | Identity | Gaps |
| --- | --- | --- | --- | --- | --- |
|  | Ch. | Range | Value | Ratio |  |
| 3B41 | 2 | 856636 to 859505 | 0.0 | 2330/2886 | 39 |
| 3B47 | 2 | 520404 to 522396 | 0.0 | 1449/2015 | 43 |

Chromosome 2 NCBI ID CP022603.1

### Clones Matching to Mycoplasma Bacteria

There are ten 3B series clones with sequences that best match to separate sequences within both *Mycoplasma fermentans* and *Mycoplasma conjunctivae* strain NCTC10147. The complete genome of this later species of mycoplasma has yet to be fully assembled. The matching stealth virus clones are listed in Table 4 in the order of their sequence alignments with increasing numbers of the matching M*. fermentans* nucleotides. For one of the partially sequenced clones (3B680), while its T7 sequence matched closely to *M. fermentans* and *M conjunctivae*, 79 of the first 85 of 100 nucleotides at its T3 end matched to a sequence in SCMV. The matching to SCMV strain 2715 (NCBI accession number FJ483968.2) extended from nucleotide 202627 to 202721 of the SCMV genome with an Expect Value of only 2e-25 due in part to its small size. The 85 nucleotides in the T3 sequence of clone 3B680 also matched to 85 nucleotides within two clones 3B418 and 3B625 that matched at both of their ends with sequences of SCMV. The overall level of matching of the sequences of the ten clones with the sequences of both *M. fermentans* and *M. conjunctivae* is 97.5% (13,931/14,287 nucleotides)

**Table 4.** Clones With Sequences That Best Match to *Mycoplasma fermentans* M64 Strain.

| Clone | Matching Nucleotides | Expect Value | Identity Ratio | Gaps |
| --- | --- | --- | --- | --- |
| 3B627 | 152708 to 153034 | 4e-163 | 327/328 | 0 |
| 3B632 | 174189 to 175572 | 0.0 | 1380/1384 | 1 |
| 3B520 | 473083 to 475878 | 0.0 | 2771/2797 | 2 |
| 3B528 | 651388 to 653431 | 0.0 | 2020/2044 | 1 |
| 3B512 | 668449 to 670792 | 0.0 | 2342/2345 | 1 |
| 3B35 | 715735 to 717874 | 0.0 | 2136/2142 | 2 |
| 3B37 T7 | 763908 to 764569 | 0.0 | 499/500* | 0 |
| T3 | 765106 to 765768 | 0.0 | 549/550* | 8 |
| 3B545 | 801182 to 801394 | 0.0 | 212/213 | 1 |
| 3B622 T3 | 962497 to 963182 | 0.0 | 587/604* | 1 |
| T7 | 963536 to 964273 | 0.0 | 607/625* | 3 |
| 3B680 T7<sup>#</sup> | 1106962 to 1107081 | 1e-49 | 118/120 | 0 |

\*There are several ambiguous sequences, designated N, in these clones. The identity ratios were determined on the sequences that preceded the first ambiguous nucleotide.
<sup>#</sup> 3B680 T3 has 79 of 85 nucleotides that match a sequence in SCMV

### Clones Preferably Matching to Other Bacteria

The following are the best currently available alignments of the sequences in some of the remaining clones. For example, there are three clones (C16122, C16127, and C16141) in which with DNA sequences best match to different species of *microbacterium* bacteria. These are small, gram-positive bacteria with a high guanosine-cytosine content that are classified within the <u>Actinomycetia</u> class of bacteria [20] . The matching of the T3 and T7 sequences from clone C16122 was to different species of this bacteria. The T3 sequence matching clone C16122 T7 was reasonably high with an Expect Value of 2e-102. However, it was only the region between nucleotides 162 to 505 of the 812 long nucleotide sequence of the clone that matched to a *microbacterium* sequence. The regions of C16122 T7 from nucleotide 1 to 161 and 505-812 showed no matching with any of the accessible GenBank sequences, even when using the BLASTX program. The e-120 and e-97 Expect Values, respectively, of the sequence alignments for clones C16127 and C16141 to microbacterium species of bacteria are reasonable in view of the relatively short lengths of the sequences. The data on the three clones are recorded in Table 4.

The T3 and T7 sequences of clone (C16125) show preferred matching to a 157689 base pair (bp) plasmid isolated from the MDW-2 unclassified species of Aminobacter bacteria (Table 5). Aminobacter bacteria are α-proteobacteria in the same *Hyphomicrobiales* Order as the *Brucella – Onchrobactrum* bacteria but belong to a different family. One clone (C16135) has a T3 sequence that best matches to a sequence within a strain of *Cellulosimicrobium cellulans* bacteria, which is also within the <u>Actinomycetia</u> class of bacteria [20]. The T7 sequence of clone C16135, however, matches to an intron sequence within the Karzin coding human cellular gene (NCBI Accession no. NG_029844.2) with 205/396 nucleotide identity after discounting the non-assigned “N” nucleotides.

**Table 5.** Clones With Sequences That Best Match to Different Species of *Microbacterium*.

| Clone | Species | Matching Nucleotides<br>Range | Expect<br>Value | Identity<br>Ratio | Gaps |
| --- | --- | --- | --- | --- | --- |
| C16122 T3 | A | 630149 to 630770 | 2e-102 | 461/624 | 2 |
| C61122 T7 | B | 1690217 to 1690563 | 6e-26 | 241/348 | 5 |
| C16127 | C | 1364049 to 1364299 | 2e-120 | 249/251 | 0 |
| C16141 | D | 657325 to 657522 | 7e-93 | 197/198 | 0 |

NCBI ID of best matching *Microbacterium* sequence and name of species
A CP063814.1 *Microbacterium* A18JL200 chromosome
B CP043732.1 *Microbacterium esteraromaticum* strain B24
C LR880474.1 *Microbacterium* Nx66 genome assembly
D CP080491.1 *Microbacterium* Se5.02b chromosome

**Table 6.** Clone With Sequences That Best Match to A Plasmid Sequence in *Aminobacter*.

| Clone | Matching Nucleotides<br>Range | Expect<br>Value | Identity<br>Ratio | Gaps |
| --- | --- | --- | --- | --- |
| C16125 T7 | 19341 to 19987 | 0.0 | 565/659 | 12 |
| C16125 T3 | 21377 to 22070 | 0.0 | 578/698 | 5 |

NCBI ID: CP060199.1 *Aminobacter* sp. MDW-2 plasmid pMDW2

**Table 7.** Clone With Sequence at One End That Best Match to *Cellulosmicrobian* strain ORNL-0100 chromosome.

| **Clone** | **Matching Nucleotides**<br>**Range** | **Expect**<br>**Value** | **Identity**<br>**Ratio** | **Gaps** |
| --- | --- | --- | --- | --- |
| C16135 T3 | 1323875 to 1324367 | 0.0 | 477/493 | 0 |

NCBI ID: CP072387.1
The T7 sequence of clone C16135 matches to a sequence in the human Karzin gene

Clone 3B513 (NCBI Accession no. U27894.2) is a fully sequenced clone of 8,106 nucleotides. Routine BLASTN analysis only identifies five regions with a matching score of >200 to bacterial sequences. Collectively, these regions comprise only 2,346 nucleotides. Moreover, only one of the matches had an Expect Value of 0.0. Far more extensive sequence matching was obtained using the BLASTX and pairwise matching of the sequence of 3B513 with the nucleotides coding the best-matching amino acid sequence. The results of this analysis are summarized in Table 8. While there was still incomplete matching for the first 2,262 nucleotides, Expect scores of 0.0 were obtained for the remaining 5,844 nucleotides. Of the three matching sequences, those matching to an *Ochrobactrum sp. POC9* sequence showed only 2 of 399 nucleotide differences to the overlapping sequence at the 5’ of a *Brucella* gallinfaecis gene and only 1 of 374 nucleotide differences to the overlapping at the 3’ end to a Mesorhizobium *denitficams* gene.

**Table 8.** Bacterial Nucleotides Matching to the Sequence of Clone 3B513.

| 3B513<br>nucleotides | Matching<br>Bacteria | Accession No. of Bacterial<br>Sequence with Matching<br>Nucleotides | Numbering of the<br>Matching Bacteria<br>Nucleotide Sequence | BLASTN Matching |  |  |  |
| --- | --- | --- | --- | --- | --- | --- | --- |
|  |  |  |  | Score | Expect | Identities | Gaps |
| 1-376<br>1212-1616<br>1913-2284<br>2399-2615 | Aliihoeflea sp.* | NZ_JAAAMH010000008.1 | 170669-171044<br>171895-172298<br>172632-172963<br>173078-173294 | 232<br>120<br>83.3<br>43.7 | 2e-63<br>7e-30<br>2e-18<br>2e-06 | 279/377<br>278/409<br>220/335<br>142/219 | 2<br>9<br>4<br>6 |
| 2663-5600 | Brucella<br>gallinifacis** | NZ_JAQNAJ010000019.1 | 1-2938 | 679 | 0.0 | 2927/2938 | 0 |
| 5202-7509 | Ochrobactrum sp<br>POC9*** | NZ_QGST01000056.1 | 284-2598 | 4108 | 0.0 | 2298/2315 | 7 |
| 7143-8106 | Mesorhizobium<br>denitrificans**** | NZ_QURN01000017.1 | 77672-78643 | 1707 | 0.0 | 962/972 | 8<sup>#</sup> |

\*Aliihoeflea sp. 40Bstr573 NODE\_8, whole genome shotgun sequence. The next highest matchings were to three separated regions of *Shinella sumterensis* (Expect values 4e-57, 2e-35, 2e-15)
\*\* *Brucella gallinifacis* strain IPG27 LOCAPJJO\_19, whole genome shotgun sequence
\*\*\* *Ochrobactrum* sp. POC9 *Ochrobactrum*\_sp.\_POC9\_contig00056, whole genome shotgun sequence
\*\*\*\* *Mesorhizobium denitrificans* strain LA-28 contig\_17, whole genome shotgun sequence
<sup>#</sup> The 8 gaps were contiguous

BLASTN analysis of clone 3B513 also revealed 94% nucleotide matching, dismissing non-assigned “N” nucleotides, with a 503 nucleotides region within the 3B525 T3 sequence extending from nucleotides 76 to 593. The matching region was within the section of the 3B513 clone that corresponded to the sequence of the *Mesorhizobium denitrificans* gene. Nucleotides 15 to 82 on clone 3B525 T3 match to SCMV (NCBI Accession FJ483968.2 with an Expect Value of 2e-09 and 52/64 identical nucleotides after omitting 4 non-assigned “N” nucleotides. The matching SCMV nucleotides were from 72,686 to 72,619. The sequence of clone 3B525 T7 matches to SCMV nucleotides 66002 to 66651 with an Expect score of 0.0. Moreover, the 3B527 T7 sequences closely matches the sequences clones 3B526T7, 3B550 T7, 3B544 T7, 3B320 T3, 3B642 T7, 3B314 T3, 3B663 T3 all of which have SCMV-related sequences in both their T3 and T7 sequences (data not shown).

## Discussion

The presence of bacterial sequences in the cultures of a stealth adapted virus cannot be explained by bacterial contamination of the cultures. This possibility is excluded by the following observations: i) Repeated blood cultures from the patient over a 4-year period gave very comparable results in terms of the observed formation of foamy vacuolated cells with prominent syncytia. ii) There were no indications for bacteria being present in the cultures throughout the frequent viewing of living cells by phase contrast microscopy, repeated examinations of hematoxylin and eosin-stained cells by regular microscopy, and studies using electron microscopy. iii) Some of the cultures were maintained in antibiotic free medium for extended time periods. iv) The identified sequences are not from the bacteria that are typically involved in bacterial contamination of tissue cultures. v) The agarose gel of the nucleic acids extracted from the pelleted material obtained from filtered and ultracentrifuged tissue culture medium shows only minimal DNA that is larger than the approximately 20 kb band and, which would have been expected if there were bacterial chromosomal DNA. vi) The C16 series of clones were derived solely from the DNA that banded in the agarose gel at a size of approximately 20 kb.

The relocation or transposition of genetic sequences from their bacterial origin to a virus can be viewed as a passive hijacking of the sequences by the virus and the loss of the bacteria’s capacity to restrain their own genetic sequences from deserting and moving elsewhere. To help in understanding this process, the relocating sequences are referred to as “renegade sequences” [9]. The term “viteria” was also introduced to describe viruses with incorporated bacteria-derived genetic sequences [8].

Even though the double stranded DNA pelleted material migrated in agarose gel with an approximate size of 20 kb, the sum of the previously reported SCMV-derived nucleotide sequences (∼100,000), cellular-derived sequences (∼ 25,000) and the presently reported bacterial related sequences (∼ 50,000) indicates that the entire genome comprises genetically different segments with a diversity of viral, cellular, and bacterial-derived sequences. These segments are not necessarily all packaged into each virus particle, which did appear to be morphologically heterogeneous on electron microscopy [1]. It is noteworthy that the agarose gel of the nucleic acids extracted from the pelleted material showed a substantial amount of RNA (Figure 1). A 20 kb size of the double-stranded DNA is consistent with RNA being involved in the replication process. Thus, 20 kb is close to the upper size limit of replicating RNA without the inclusion of an intrinsic genomic proofreading mechanism [21]. Involvement of RNA may also explain the genetic instability previously reported in the SCMV and cellular related DNA genetic sequences [3]. It is also consistent with a reverse transcriptase step being required to obtain a positive PCR in a stealth adapted virus culture from a different CFS patient [22].

The mechanism of the apparent incorporation of bacteria-derived sequences is probably the same as that occurring with the cellular-derived sequences in this and in other stealth adapted viruses [9,10]. It is perceived as single stranded RNA cross-linking residual DNA or RNA segments remaining from the originating SCMV virus after it has undergone fragmentation as part of the stealth adaptation process. Further fragmentation of the virus could lead to additionally incorporated cross-linked genetic sequences. The finding of bacterial and SCMV related sequences at the opposite terminal regions of clones C16282, 3B680, and 3B525 is consistent with this hypothesis. This possibility is further supported by the limited SCMV matching of nucleotides 15 to 82 on clone 3B525. This region slightly overlaps with the bacterial matching sequences from nucleotides 76 to 593 in the same clone. Also consistent with a cross-linking process is the presence of a bacterial sequence and a cellular sequence at the T3 and T7 readouts, respectively, of clone C16135.

Another example of overlapping sequences is seen within clone 3B513. The end regions of the *Ochrobactrum*-matching sequences in clone 3B513 overlap with the end regions of Brucella gallinifaecis and of *Mesorhizobium denitrificans* bacterial sequences, respectively. Moreover, as mentioned above, a region within the *Mesorhizobium*-related sequence matches to a region within clone 3B525 T3. In addition to sequences linking to each other, homologous recombination can lead to the substitution of sequences with matching end regions (9). It is still possible, however, that the predominant viral, bacterial, and cellular sequences are mainly located on discrete segments of the stealth adapted virus. This question can be resolved by further sequencing of the virus, which is archived at the American Type Culture Collection (ATCC).

A subsequent article will discuss the potential functions of the incorporated bacterial genetic sequences in more detail. At least some of the coded proteins presumably contribute to the virus’s replication and transmission. One such role could be providing alternative or additive capsid-like proteins [13]. Some of the proteins are enzymes and have potentially useful metabolic functions. A self-healing process occurs during the culturing of the stealth adapted viruses [23]. It is associated with the production of self-assembling aliphatic and aromatic chemical compounds. The assembled materials are referred to as alternative cellular energy (ACE) pigments [23] These materials can attract an external force provisionally called KELEA (Kinetic Energy Limiting Electrostatic Attraction), which drives the ACE pathway [24]. Enhancing the ACE pathway can lead to the suppression of the virus induced CPE [24] It could be, therefore, that some of the bacteria gene-coded proteins are contributing to the formation of ACE pigments and, thereby, preventing total viral elimination of the infected cells. Similarly, intracellular protein aggregates can trigger the unfolded protein response [25] that can also delay virus-induced cell death.

An intracellular location for interactions between bacterial and viral sequences is easier to envision for those bacteria that can replicate intracellularly. This applies to infections with *mycoplasma*, *brucella,* and *microbacterium* bacteria [14, 26]. Intracellular growth in mammalian cells is not, however, a known characteristic of *Ochrobactrum* bacteria [26–28]. These bacteria are mainly viewed as being present in soils and comprising part of the complex rhizosphere surrounding and commonly penetrating plant cells. Indeed, considerable symbiosis occurs between plant cells and a wide array of endophytic bacteria [29]. Another option is for stealth adapted viruses to directly enter soil-based bacteria, which can be present in consumed uncooked foods [30]. Some of these ingested bacteria may continue to reside in the gut microbiota. Interestingly, atypical bacteria were cultured from the feces of the CFS patient from which stealth virus-1 was cultured. Moreover, transmissible cytopathic activity was subsequently retrieved from the atypical bacterial colonies (unpublished). There are major epidemiological ramifications if bacteria can be involved in the transmission of stealth adapted viruses.

A related characteristic of stealth virus-1 is its wide host range, including being infectious for insect cells [2] . This could occur through the deletion of genes responsible for the typical species restricted growth of most animal cytomegaloviruses. Insects also commonly harbor endophytic bacteria [31].

O. *quorumnocens* strain A44 was originally isolated in Holland from the rhizosphere of field potatoes [17] . It has a defining function of metabolically inhibiting *N*-acyl homoserine lactones, a chemical used by certain gram negative bacteria to establish a more pathogenic quorum that can lead to “soft rot” [17] Because O. *quorumnocens* strain A44 lacks the 16S-28S intervening sequence present in clone 3B43, it cannot be regarded as the unequivocal source of the O. *quorumnocens* related sequences. Moreover, the *B. pseudogrignonensis* related sequences in two of the clones (3B41 and 3B47) are not present in O. *quorumnocens;* nor are the *O. quorumnocens-*related sequences in clone 3B513 directly identifiable with strain A44.

Intact B*. pseudogrignonensis* bacteria are noteworthy because of their ability to induce tumors in mushrooms [32]. This is mentioned to underscore the uncertainty as to the potential biological consequences of the virus mediated transfer into humans of infectious bacteria-derived genetic sequences. Moreover, the exact origins and biological functions of the incorporated bacterial sequences may remain in doubt because of their genetic instability with potential ongoing divergence from the original bacterial sequence. As seen with incorporated cellular sequences, there is also the potential for the substitution between different bacterial sequences [10]. This is most apparent in clone 3B513.

As noted above, it is easier to envision the assimilation of *mycoplasma* sequences into stealth adapted viruses than from soil-based bacteria. It is particularly noteworthy that the closest alignment is with *M. fermentans,* strain M64. (There was essentially identical nucleotide matching to *M. conjunctivae*, a sheep eye pathogen [33].) The *M. fermentans* species of mycoplasma gained special interest during the early history of the AIDS epidemic. Although HIV had been isolated from patients, concerns were expressed that a co-infecting pathogen was also required for the development of severe illness. Dr. Shyh-Ching Lo identified anti-mycoplasma antibodies and mycoplasma DNA sequences in several AIDS patients with more fulminant disease. He was eventually able to isolate a culturable mycoplasma, which is called M. incognitus. It was later confirmed as *M. fermentans incognitus* [34–38].

Most of the HIV clinical studies, however, relied on serological and molecular testing rather than on isolating culturable bacteria [34–38]. Indeed, the isolation of an intact *mycoplasma* bacteria could be more of a coincidental and somewhat misleading distraction from finding a stealth adapted virus with incorporated *M*. *fermantans* genetic sequences.

The clinical association between positive *M. fermentans* assays and the severity of HIV illness is relevant to the proposed role that the testing of an experimental polio vaccine in chimpanzees had in the formation of HIV [39]. The polio vaccine was grown in Rhesus monkey kidney cell cultures. It was reportedly contaminated with a cytopathic virus that was difficult to culture [40]. It also has detectable DNA of rhesus monkey cytomegalovirus [41]. These two findings are consistent with the vaccine being contaminated with a stealth adapted virus able to promote the growth and transformation of simian to human immunodeficiency virus.

Several other illnesses, including CFS, Fibromyalgia, Gulf War Syndrome, and chronic arthritis were previously attributed to ongoing infection with *M. fermentans* bacteria [42–46]. This conclusion was based mainly on positive serology and molecular detection methods. Clinical trials with long term antibiotic therapy failed to achieve clinical remission and the presumed bacterial infections were largely dismissed as being coincidental to the real cause. Based on the data presented in this article, the findings are consistent with a viteria infection, in which the underlying stealth adapted virus has incorporated mycoplasma-derived genetic sequences. An interesting admission by the owner of a commercial mycoplasma molecular testing facility for both CFS and Gulf War Syndrome patients was that the patient’s PCR amplified products commonly showed minor genetic differences. Although he reported the results as being positive for *M. fermentans,* he inwardly thought the patients were infected with a multiplicity of mycoplasma strains, none of which had the exact sequence of *M. fermentans*.

There are also published studies suggesting the potential involvement of *Brucella* bacteria in CFS, fibromyalgia, and the Gulf War Syndrome [47]. Again, this conclusion was primarily based on positive serological and molecular detection methods. This was also the suggestion of an inadvertent consequence of efforts to use *Brucella* bacteria to create a Germ Warfare agent [48]. The data provided in this article are more consistent with infection with a viteria containing *brucella* bacteria-derived genetic sequences.

CFS is an imprecisely defined illness [49] . It can range in severity from a rather mild illness to the patients having severe cognitive impairments. CFS has many clinical features in common with another somewhat controversial illness termed chronic Lyme disease [50–55]. This supposedly tick-borne illness is attributed to chronic infection with *Borrelia burgdorferi* bacteria. Comparable to the methods for attributing CFS to mycoplasma infection, the evidence for Borrelia infection is largely serological and/or molecular. Many of the self-designated “Lyme Literate” physicians contend that the disease is empirically diagnosable even with negative *Borrelia* testing results. Serological and molecular studies have further suggested that many chronic Lyme disease patients are commonly coinfected with other types of bacteria, including strains of *Bartonella, Ehrlichia*, *Anaplasma*, and *Rickettsia* Bacteria [56]. Infection with *Babesia*, a unicellular parasite, is also thought to be common. Blood samples from many patients diagnosed with chronic Lyme disease were positive when personally tested for stealth adapted virus infection (unpublished data). It will be of interest to sequence stealth adapted viruses cultured from patients diagnosed as having chronic Lyme disease. *Borrelia* infections are increasingly being linked with other illnesses. These include Morgellons disease [57], in which the patients have skin lesions from which electrostatic particles can be obtained. Other diseases attributed to *Borrelia* include acute psychosis, carditis, autoimmunity, and Guillain-Barre Syndrome [58].

Similar considerations apply to other illnesses that are likely to be due to infection with stealth adapted viruses but are publicly being mistakenly as a bacterial infection. These illnesses include PANDAS (Pediatric Autoimmune Neuropsychiatric Disorders Associated with Streptococcal Infections) [59–61]. Blood samples from several children with this diagnosis were tested for stealth adapted viruses with positive results (unpublished).

Other bacterial infections have presumptively been associated with a range of additional neurological, psychiatric, dermatological, and allergic disorders [62–66]. Consideration of viteria, i.e., stealth adapted viruses with incorporated bacteria-derived genetic sequences, may help reorient thinking about these presumed associations.

Regardless of the presence of bacteria-derived genetic sequences, major consideration should be given to the demonstrated wide range of neurological, psychiatric, and other illnesses caused by stealth adapted viruses. Positive viral cultures were regularly obtained from blood samples of autistic children [67] and from children with severe learning and behavioral disorders [68]. A virus inducing a very similar CPE as did stealth virus-1 was isolated from the cerebrospinal fluid (CSF) of a comatose patient with a four-year history of bipolar psychosis [69]. Multiple family members have received different diagnoses and yet have experienced similar core symptoms. The diagnoses in one family were dementia in both grandparents, CFS in their daughter, amyotrophic lateral sclerosis (ALS) in her husband, and major learning disorders in three younger children [70]. A controlled study provided positive culture results in all 10 tested patients with multiple myeloma [71]. Animal illnesses can also be caused by stealth adapted viruses [72]. On a more positive note, evidence of recovery from stealth adapted virus infections highlights the role of the ACE pathway as a non-immunological anti-viral defense mechanism [23, 73].

In summary, this article details the approximate derivations of the bacteria-derived sequences present in the culture of a stealth adapted African green monkey simian cytomegalovirus (SCMV) infecting a patient with chronic fatigue syndrome. Most of the bacterial sequences are related to those of either *Ochrobactrum* or *Mycoplasma*. These and additional bacterial sequences were seemingly derived by genetic recombination. The data are relevant to the biology of stealth adapted viruses. The data also can explain the potential of misdiagnosing a stealth adapted virus infection as a bacterial infection. Additional DNA sequencing of cultured stealth adapted viruses is indicated.

## Abbreviations

ACE – Alternative Cellular Energy, CFS – Chronic Fatigue Syndrome, CPE – cytopathic effect, CSF – cerebrospinal fluid, KELEA – Kinetic Energy Limiting Electrostatic Attraction, PCR – Polymerase Chain Reaction, SCMV – African green monkey simian cytomegalovirus, AIDS – Acquired Immunodeficiency Disease, N - non-assigned nucleotide, nt – nucleotide, bp – base pair, kb – kilobase

## Acknowledgement

Research on stealth adapted viruses is supported by MI Hope Inc., a non-profit public charity based in South Pasadena CA.

